# What are the demographic consequences of a seed bank stage for columnar cacti?

**DOI:** 10.1101/2020.11.24.396374

**Authors:** Gabriel Arroyo-Cosultchi, María C. Mandujano, Roberto Salguero-Gómez, Armando J. Martínez, Jordan Golubov

## Abstract

The dynamics of plants populations are often limited by the early stages in their life cycles. The question if the columnar cacti have or not a seed bank in predictable environments. Yet, information regarding seed bank dynamics and how these may influence the full life cycle of plant species is remarkably scarce or ignore. This lack of knowledge is mostly due to the challenges in quantifying seed vital rates. Studies of arid land plant species have historically been focused on the drivers of sporadic recruitment. However, little attention has been given to the demographic consequences of early developmental stages, including seed banks. Here, we evaluate the effects of seed bank survival and seedling recruitment vital rates on the population dynamics and viability of 12 columnar cacti species, recent evidence suggests that cacti seeds may remain viable for the short-term. We assess how changes in the vital rates of these processes, and the inclusion of a seed bank affect population growth rate (*λ*). We found that a seed bank in the examined matrix population models significantly increased *λ* as well as the vital rate elasticities of *λ* to growth and fecundity, whereas that of overall survival decreased. Our numerical simulations showed that seed survival had a larger effect on *λ* than seedling recruitment and establishment. We suggest that seed bank may explain the structure and population dynamics. Thus, we argue reconsider that this early stage in demographic models will generate more informed decisions on the conservation and management of columnar cacti.

## 1 INTRODUCTION

In plant populations, seeds and seedlings often act as primary constraints to population size and demographic viability (Ågren, 1996; Crawley, 1990), particularly true for short to mid-lived species (Silvertown et al., 1993; Franco and Silvertown, 2004). On the one hand, seed limitation whereby the number of individuals increases following seed addition (Turnbull et al., 2000), occurs when insufficient amounts of seeds are produced (Ågren, 1996), when these are not viable (Bell et al., 1993) or when their dispersal is limited (Clark et al., 2007). On the other hand, the establishment of those seeds as seedlings may be limited by factors such as environmental stress, cross-pollination, pollinator limitation (Bell et al., 1993) or microsite availability (Eriksson and Ehrlén, 1992; Turnbull et al., 2000), regardless of seed limitation (Clark et al., 2007). Seedling recruitment is understood as the process by which new individuals are added to a population, including seed germination, seedling survivorship, and seedling growth (Eriksson and Ehrlén, 2008).

Seedling recruitment in cactus species is largely recognized as a critical process in their life cycles, and thus as an important restriction to population growth rate *λ* (Mandujano et al., 1996; Godínez-Álvarez et al., 2003; Martínez et al., 2010; Arroyo-Cosultchi et al., 2016). While several studies have examined the mechanisms limiting recruitment in the Cactaceae, these have predominantly assessed establishment limitation (Steenbergh and Lowe, 1977; Cody, 1993; Godínez-Álvarez et al., 2003; Mandujano et al., 2007; Holland and Molina-Freaner, 2013). However, the demographic consequences of seed and seedling limitation in this diverse taxon have been largely overlooked. Studies in Cactaceae show a high potential germination rate in laboratory conditions, which contrasts with the high seedling mortality rates reported in the field (Esparza-Olguín et al. 2002; Pierson et al. 2013; Holland and Molina-Freaner 2013; Zepeda-Martínez et al. 2013). Together, this group of research suggests that demographic processes at the interface of seed limitation and seedling recruitment are crucial to life cycles and to population dynamics in Cactaceae.

An important trait for plant species inhabiting unpredictable environments such as arid lands, is the ability to generate seed banks (Gutterman, 1994). This strategy is thought to be a fundamental component for population persistence in variable environments (Pake and Venable, 1996) and it is, therefore, a key factor that affects seed and seedling limitation (Venable, 2007). However, estimating seed bank dynamics is challenging as seed longevity depends on morphological and physiological traits such as seed size, dormancy, and photoblastism (Baskin and Baskin, 1989; Rojas-Aréchiga and Batis, 2001; Rojas-Aréchiga, 2014), as well as on mortality by biotic drivers such granivores and pathogens (Álvarez-Espino et al., 2014). So obtaining accurate estimates for survival and germination of seeds (often times of a more few millimeters) in the soil can be an arduous task (Adams et al., 2005; Nguyen et al., 2019).

Even though the presence/absence of a seed bank can be important in demographic terms (Kalisz and McPeek, 1992; Doak et al., 2002), these are not always included in plant demographic models (Nguyen et al., 2019). This is so albeit evidence of their presence in numerous species (Doak et al., 2002; Nguyen et al., 2019). Still, this life stage is commonly assumed to be non-existent or short-lived or the origin of seedlings is not differentiated. Seeds residence time in the soil determines if the species may generate a transient (<1 year), short-term (≥ 1 but <5 years) or long-term persistent seed bank (≥5 years) sensu (Bakker et al., 1996; Thompson et al., 1997). The presence of seed banks is often correlated with a short mean life expectancy of established individuals (Cohen, 1966); however, seeds dormancy and the formation of a seed bank are potentially costly features (Rees, 1994). Additionally, seed banking in long-lived organisms can serve as a hedge against recruitment failure from periodical fluctuations in seed production and/or seedling recruitment (Rees, 1994).

The inclusion - or not - of seed banks in population models can affect the assessment of the population dynamics (Doak et al., 2002; Nguyen et al., 2019). In existing population models in the Cactaceae, seed banks are rarely explicitly considered (Schmalzel et al., 1995; Godínez-Álvarez et al., 2003; Zepeda-Martínez et al., 2013; Mandujano et al., 2015). Indeed across columnar cacti, most demographic studies do not include seed banks (Godínez-Alvarez et al., 1999; Esparza-Olguín et al., 2005; Rojas-Sandoval and Meléndez-Ackerman, 2013), except in Cephalocereus polylophus (Arroyo-Cosultchi et al., 2016). Assuming no seed banks in the Cactaceae may be correct if seed viablitiy quickly decreases after dispersal (seed limitation; Rojas-Aréchiga and Batis, 2001; Méndez et al., 2004), or if granivore pressure is high (dispersal limitation; Valiente-Banuet and Ezcurra, 1991; Sosa and Fleming, 2002; Clark-Tapia et al., 2005). However, growing evidence suggests that cactus seeds may remain viable for significant periods of time (e.g. 1-2 years) (Mandujano et al., 1997), leading to a potential short-term seed bank (Godínez-Álvarez et al., 2003; Bowers, 2005; Cano-Salgado et al., 2012; Álvarez-Espino et al., 2014; Ordoñez Salanueva et al., 2017; Lindow-López et al., 2018).

Matrix population models are useful tools in plant population ecology, as they provide a common conceptual framework for comparative research (Silvertown et al., 1993; Salguero-Gómez and de Kroon, 2010; Salguero-Gómez and Plotkin, 2010; Nguyen et al., 2019). The growing number of studies using comparative approaches with matrix population models (Salguero-Gómez et al., 2015) has allowed linking specific vital rates (Franco and Silvertown, 2004; Adier et al., 2014) and stages in ecological successional gradients (Silvertown et al., 2002), life history evolution (Burns et al., 2010), population dynamics of native *vs*. invasive plant species (Ramula et al., 2008), or senescence (Baudisch et al., 2013; Jones et al., 2014) among others. We focused on the vital rates of seed bank survival and seedling recruitment to examine their relative effects on the overall population growth rates (*λ*). We apply matrix population models to assess (i) the role of seed banks and related dynamics (e.g. recruitment of seedling from seed bank) as opposed to recruitment from direct reproduction, *i*.*e*. without seed banks, on the vital rate elasticities and population growth rate (*λ*) and, (ii) evaluate the potential effects on *λ* of an increase in the vital rate of seed banks, the seed-seedling transition and seedling survival on *λ*.

## 2 MATERIALS AND METHODS

### 2.1 The Database

We used a comparative approach to determine the effects of a seed bank stage on the population dynamics of columnar cacti using published matrix population models. We searched the ISI Web of Science and Scopus electronic databases using the keywords *“columnar cacti”, “demography”, “population model”*, and *“population growth rate”* since September 1993. We included studies that explicitly used a matrix population model for columnar cacti belonging to the taxonomic tribe Pachycereeae and Trichocereeae (Anderson, 2001). Additional studies were obtained by studying the latest issues of ecological journals and by including data from (http://www.dgbiblio.unam.mx/index.php/catalogos Accessed 30 July 2015), the COMPADRE Plant Matrix Database (Salguero-Gómez et al., 2015, see Table 1), as well as part of collective, ongoing unpublished research. Our criteria for study selection included at least one matrix population model (Caswell, 2001) to estimate the population growth rate (*λ*). If the study had more matrices (*i*.*e*.>1 annual transition or populations), matrices were averaged across years or sites to obtain a single, representative matrix model per species (see Supplementary Appendix A for all original mean matrices). The final sample size contained 12 matrices, one matrix for each columnar cacti species (Table 1).

**TABLE 1.**
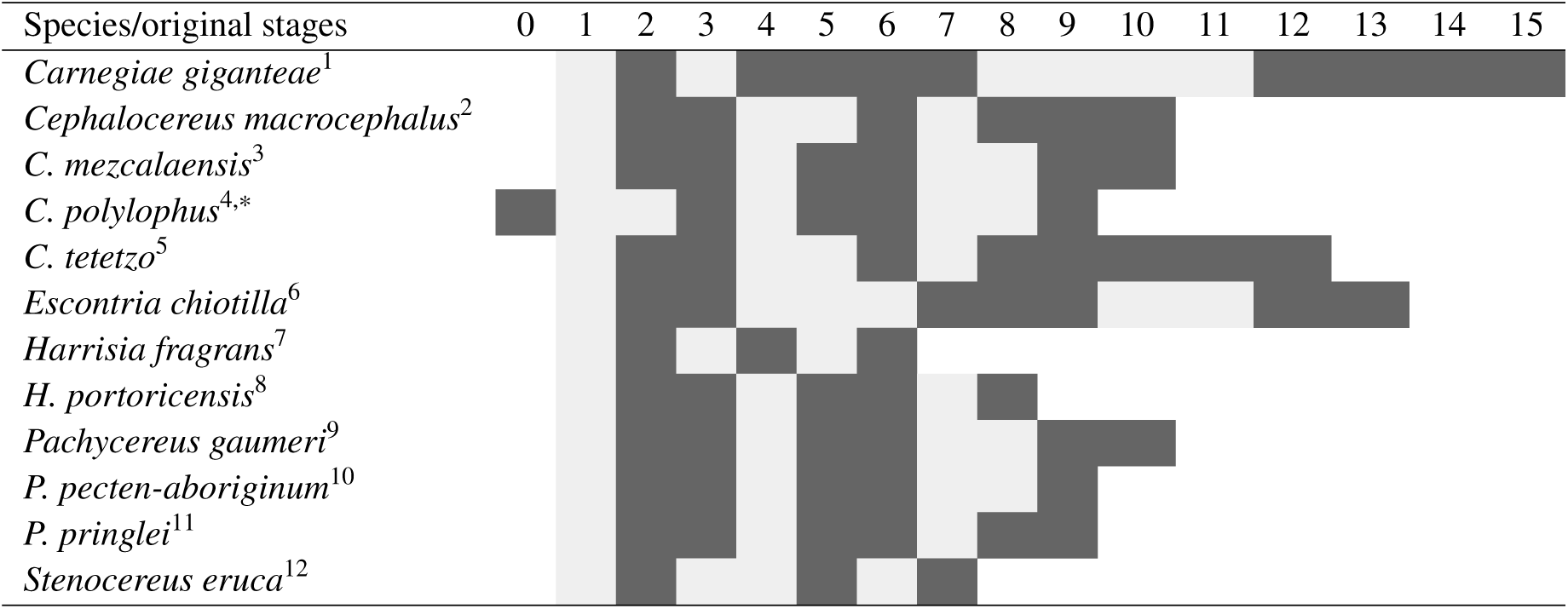
Species used in this study for which matrix population models are available, showing the original dimension of the matrices (in gray), and the adjacent life cycle classes that were reduced (in black) to produce matrices with the same dimension: 6 × 6. Stage 1 was always kept unaltered because it contains the seedling stage. *Note that all matrices lack seed bank (stage 0), except for *Cephalocereus polylophus* (stage 0, Figure 1 b). ^1^Silvertown et al. (1993), ^2^Esparza-Olguín et al. (2002), ^2,3,5^Esparza-Olguín et al. (2005), ^2,5^Godínez-Alvarez and Valiente-Banuet (2004), ^4^(Arroyo-Cosultchi et al., 2016),^5^Godínez-Alvarez et al. (1999), ^6^Ortega (2001),^7^Rae and Ebert (2002), ^8^Rojas-Sandoval and Meléndez-Ackerman (2013),^9^Méndez et al. (2004), ^10^Morales-Romero et al. (2012), ^11^Silva (1996), and ^12^Clark-Tapia et al. (2005).

### 2.2 Reduction of matrix dimensions

A critical step in comparative stage-structured demographic studies is the selection of the dimension (stage or size classes) as dimensionality affects *λ* and derived metrics (Enright et al., 1995; Ramula and Lehtilä, 2005; Salguero-Gómez and Plotkin, 2010; Picard and Liang, 2014). To overcome potential biases in our comparative inference on the role of seed bank survival and seedling recruitment for population dynamics, we standardized the variable matrix dimensions in our study, ranging originally from 15 × 15 for *Carnegiae giganteae* to 6 × 6 for *Harrisia fragrans* (Table 1). To test whether changes in matrix dimension significant changed vital rates we chose matrix dimensions of 6 × 6, 5 × 5, and 4 × 4 without a seed bank stage (hereafter WOSB) and the inclusion of a seed bank resulted in 7 × 7, 6 × 6, and 5 × 5 matrix dimensions respectively (hereafter WSB, see Supplementary Appendix A for all original and reduced matrices). We used the algorithm developed by Salguero-Gómez and Plotkin (2010) for size/stage-based matrices, adapted from Hooley (2000) for age-based models. This algorithm allows the reduction of a given matrix population model of *n* × *n* dimensions into *m* × *m*, where *m < n*. There are naturally different ways of reducing a matrix population model of interest of *n >* 2; here, we followed the recommendation by Salguero-Gómez and Plotkin (2010), whereby early life cycle stages (e.g. Figure 1) were left unaltered as they were also the life stages of interest for this study. This method preserves population growth rates, stable class distributions, and reproductive output, through the assumption of stationary stability (Salguero-Gómez and Plotkin, 2010).

**FIGURE 1.**
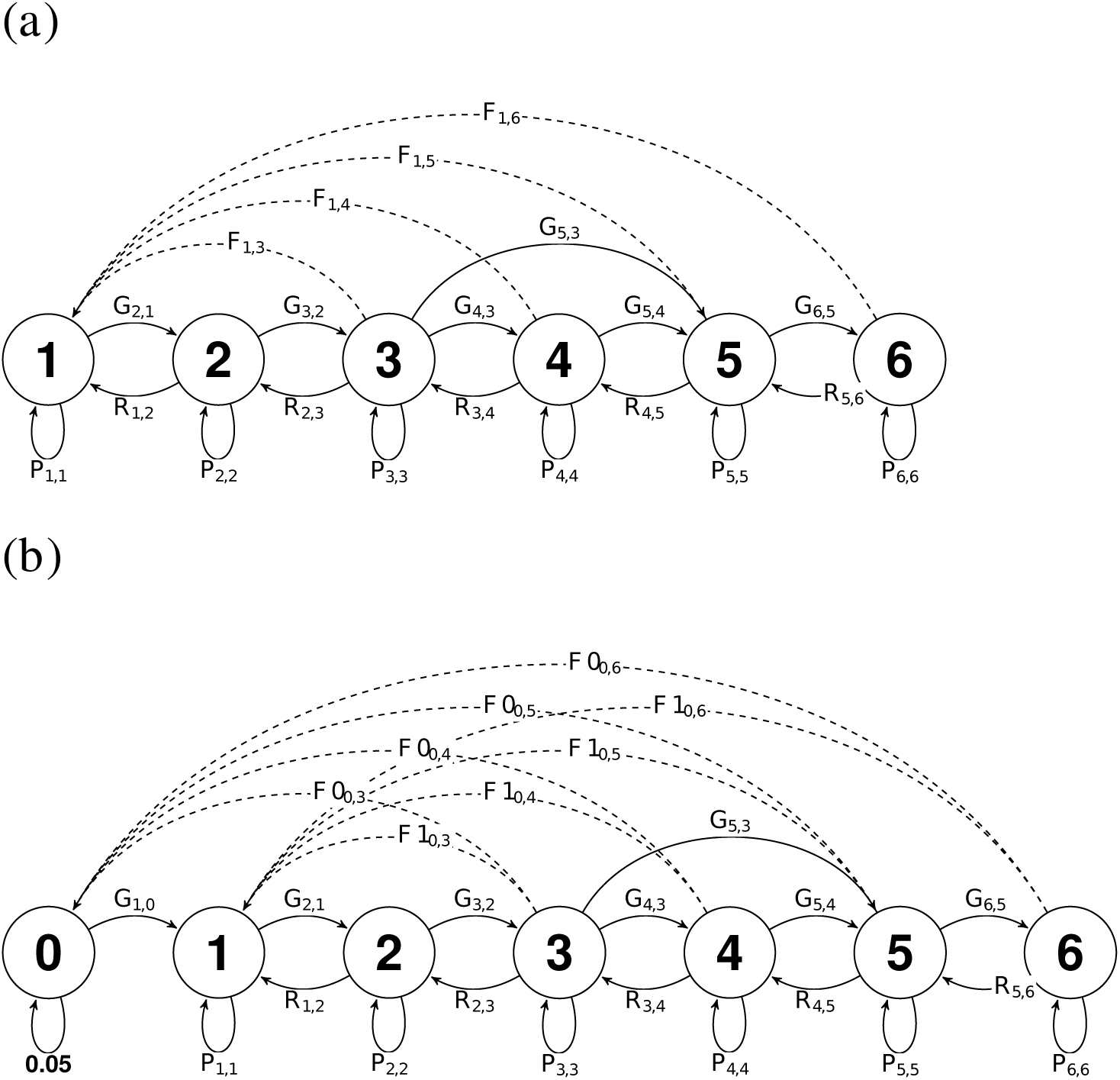
Life cycle diagrams of the matrix population models used: (a) WOSB, and (b) WSB models. Six and seven classes of individuals are possible: seeds (0), seedling (1), juvenile (2), and reproductive adults (3-6). The arrows represent the following demographic elements: stasis (P), retrogression (R), growth (G), and fecundity (F; dashed lines). The transition rate (F0) gives the fecundity into the seed bank and (F1) gives the fecundity into the seedling stage.

Matrix population models in our study were reduced by *n* - 1 dimensions by merging the two adjacent size categories with the lowest number of individuals as reported by the population vector *n(t)* while leaving the remaining stages unaltered. Here, we did not reduce reproductive and non-reproductive stages into the same class and the stage corresponding to seedlings was kept unaltered (Table 1).

The number of stages for each matrix was reduced by combining information for adjacent stages to generate new estimates of survival in a given stage class *j* (*σ*,_*j*_), negative growth (*ρ*_*ij*_), positive growth (*γ*_*ij*_), individual fecundity (*ϕ*_*ij*_), and individual ramet production (*κ*_*ij*_) (Franco and Silvertown, 2004). Fecundity entries were estimated from the information found in the original source (see Section 2.1) as the *per capita* number of seeds in each reproductive size category (Table 2). The seed to seedling transition was reported as the number of seeds × seed germination. When seed germination was calculated from laboratory and experiments under natural conditions in different sources we averaged both germination percentages as germination in natural conditions usually includes factors that affect or limit germination (granivory, drought, and fungal attack). Seedling survival was calculated by the survival of seedlings in field or laboratory conditions reported in each study (Table 2).

**TABLE 2.**
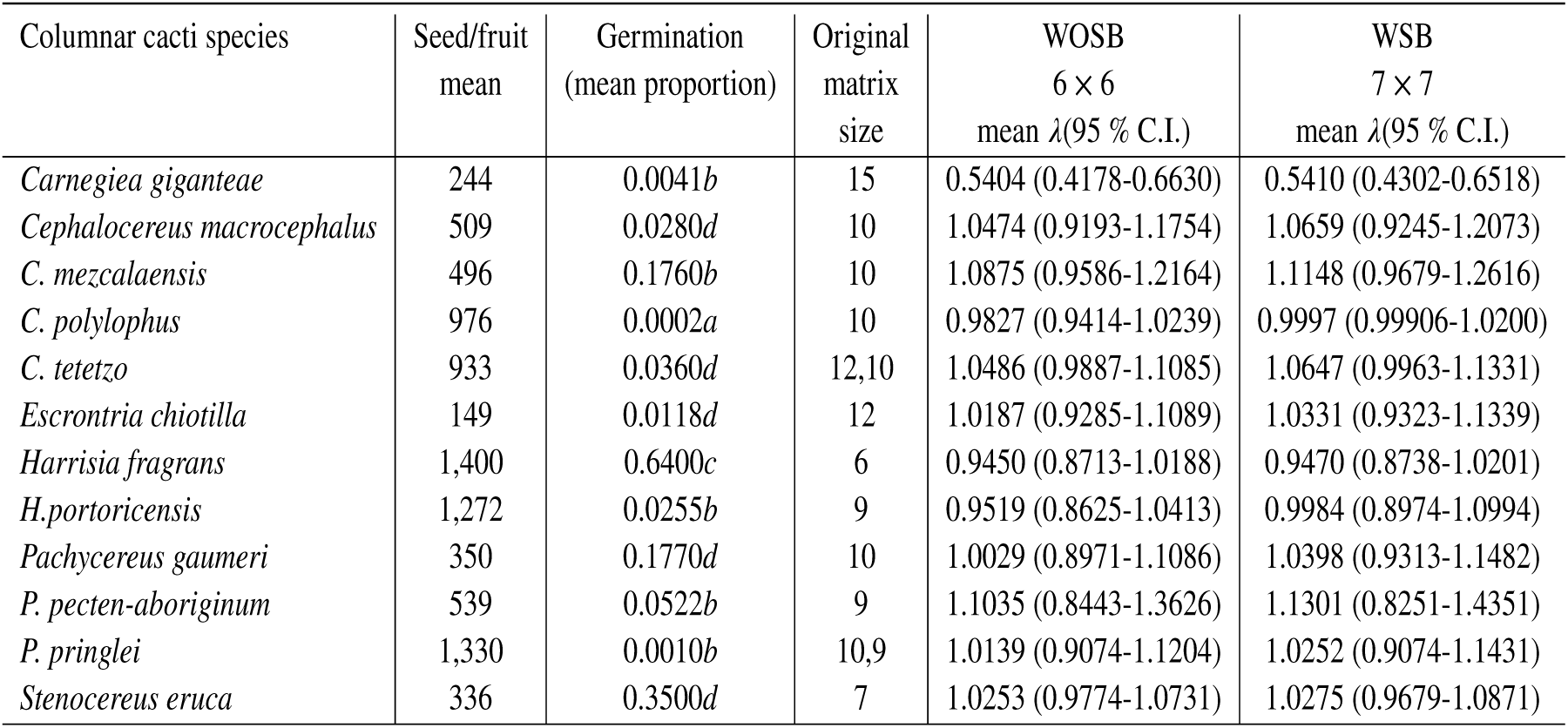
Data for the 12 columnar cacti species used in the study. Seed *per* fruit, germination (mean proportion) and matrix size corresponds to the original (no reduced dimension) matrix reported in each study, and the value of *λ* C.I. 95 % for WOSB model (6 × 6) and WSB model (7 × 7) with a hypothetical short-term seed bank (=0.05; Figure 1 b). *a* = natural *in situ* estimates, *b* = experimental *in situ* estimates, *c* = experimental *ex situ* estimates and *d* = combined *ex* and *in situ* estimates.

| Columnar cacti species | Seed/fruit mean | Germination (mean proportion) | Original matrix size | WOSB $6 \times 6$ mean $\lambda$ (95 % C.I.) | WSB $7 \times 7$ mean $\lambda$ (95 % C.I.) |
| --- | --- | --- | --- | --- | --- |
| *Carnegiea gigantea* | 244 | 0.0041*b* | 15 | 0.5404 (0.4178-0.6630) | 0.5410 (0.4302-0.6518) |
| *Cephalocereus macrocephalus* | 509 | 0.0280*d* | 10 | 1.0474 (0.9193-1.1754) | 1.0659 (0.9245-1.2073) |
| *C. mezcalaensis* | 496 | 0.1760*b* | 10 | 1.0875 (0.9586-1.2164) | 1.1148 (0.9679-1.2616) |
| *C. polylophus* | 976 | 0.0002*a* | 10 | 0.9827 (0.9414-1.0239) | 0.9997 (0.99906-1.0200) |
| *C. tetetzo* | 933 | 0.0360*d* | 12,10 | 1.0486 (0.9887-1.1085) | 1.0647 (0.9963-1.1331) |
| *Escrotria chiotilla* | 149 | 0.0118*d* | 12 | 1.0187 (0.9285-1.1089) | 1.0331 (0.9323-1.1339) |
| *Harrisia fragrans* | 1,400 | 0.6400*c* | 6 | 0.9450 (0.8713-1.0188) | 0.9470 (0.8738-1.0201) |
| *H. portoricensis* | 1,272 | 0.0255*b* | 9 | 0.9519 (0.8625-1.0413) | 0.9984 (0.8974-1.0994) |
| *Pachycereus gaumeri* | 350 | 0.1770*d* | 10 | 1.0029 (0.8971-1.1086) | 1.0398 (0.9313-1.1482) |
| *P. pecten-aboriginum* | 539 | 0.0522*b* | 9 | 1.1035 (0.8443-1.3626) | 1.1301 (0.8251-1.4351) |
| *P. pringlei* | 1,330 | 0.0010*b* | 10,9 | 1.0139 (0.9074-1.1204) | 1.0252 (0.9074-1.1431) |
| *Stenocereus eruca* | 336 | 0.3500*d* | 7 | 1.0253 (0.9774-1.0731) | 1.0275 (0.9679-1.0871) |

### 2.3 The importance of the seed bank

In the species that fulfilled our criteria, we included a hypothetical short-term seed bank (≥1 year) with an initial survival value of 0.05, except in *C. polylophus* where the transient seed bank is known (Arroyo-Cosultchi et al., 2016). In the WSB model individuals in an unstructured seed bank assumed no senescence and are thus potentially immortal. Since a seed bank is a discrete stage class and did not involve categorization, transition rates for the other classes should remain unaffected by its inclusion (Nguyen et al., 2019), however as a consequence, a temporal component in terms of longer life cycles was added. After that, the finite rate of population increase (*λ*), the stable structure of each stage (*w*), and the specific reproductive value per stage (*v*) were calculated using the WOSB (6 × 6) (Figure 1 a) or WSB (7 × 7) (Figure 1 b) models for each species (Caswell, 2001). To test for the significance of a seed bank on the population dynamics of our examined species, we used a paired *t*-test (*α* = 0.05) using the values of *λ* obtained from the WOSB and WSB models. The elasticity (*e*_*ij*_) and sensitivity (*s*_*ij*_) matrices were calculated using the *v* and *w* vectors (Caswell, 2001) and finally, 95 % confidence intervals for *λ* were estimated in order to use the analytic method suggested by Alvarez-Buylla and Slatkin (1991).

We explored the role of species, matrix dimension and seed bank on vital rates elasticities for WOSB ansd WSB models. Elasticities were calculated from each of the following vital rates: survival in a given stage class (*σ*), negative growth (*ρ*), positive growth (*γ*), and individual fecundity (*ϕ*) (Silvertown et al., 1993; Franco and Silvertown, 2004) for WOSB (6 × 6, 5 × 5, and 4 × 4) and WSB models (7 × 7, 6 × 6, and 5 × 5). We used PCA to summarize the correlation among elasticities of vital rates and included species and presences/absence of a seed bank as variables, keeping matrix dimension as a factor. PCA scores were extracted from the first four principal components (PC1 to PC4) for all variables and identified the most important variable among factor loadings. Finally, we conducted a one-way ANOVA (*α*= 0.05) on the first four principal component scores (PC1-PC4), and *post hoc* Tukey tests (*α*= 0.05) using matrix dimension as the explanatory variable. PCA’s were performed with the prcomp function of the “stats” R library (R Development Core Team, 2017).

### 2.4 Numerical simulations

The relative importance of seed bank survival and seedling recruitment from the seed bank was evaluated through numerical simulations (Adams et al., 2005; Nguyen et al., 2019). As seedling recruitment was reported in studies, the average probability of germination and fecundity were used as a proxy for the transition from seed to seedling. No clonal reproduction into the seedling stage happened such that the observed seedlings only consisted of two components: the individuals that germinated immediately between year *t* and *t* + 1 and those that germinated from the dormant seed bank from prior years (Figure 1 b). The probability of germinating within the census year is equal to the probability of germinating from the seed bank (Kalisz and McPeek, 1992).

We conducted simulation experiments to explore the influence on *λ* when vital rate probabilities during the first life stages were modified. With these simulations, we assessed the possible effects of a seed bank on columnar cacti populations during rare but potentially important events with exceptionally high or low seedling recruitment and establishment. Despite their rarity, these types of events can have substantial impacts on long-term population dynamics (Morris and Doak, 2002). The frequency and effects of these events are highly uncertain for columnar cacti, so we covered a wider range of seed survival potential, seedling recruitment and establishment probabilities and, impacts on the simulations to highlight recruitment events that are likely to be important for population growth rates. All simulations were performed independently for the WOSB (6 × 6), and WSB (7 × 7) models. We, therefore, modified the following entries depending on the presence/absence of a seed bank: seed bank (*σ*,_*sb*_), seedling survival (*σ*,_*se*_), recruitment of seedlings from the seed bank (*γ* _*sb*−*se*_), the transition from seedling to juvenile (*γ* _*se*−*ju*_), and juvenile survival (*σ*,_*ju*_). These entries of the vital rates were modified during each simulation and *λ* was calculated keeping all other vital rates constant but checking that the stage-specific survival would not exceed 1. All demographic analyses and numerical simulations were done in R (R Development Core Team, 2017) using popbio (Stubben and Milligan, 2007).

## 3 RESULTS

### 3.1 Population dynamics

Original matrices of the columnar cacti concentrated individuals in juvenile and young adult size categories except in *Cephalocereus tetetzo* and *C. polylophus*. The former had the highest proportion of individuals in the seedling stage from experimental data (no seedlings under natural conditions) so was still an approximation to naturally occurring seedlings and is very likely to be an overestimation (Godínez-Alvarez et al., 2002) and the latter quantified natural recruitment in natural conditions. *Harrisia fragrans* and *H. portoricensis* had consistently large proportions of individuals in adult size categories. Values of *λ* were not different from equilibrium for most species (Table 2); except for *Carnegiea gigantea, C. polylophus, H. fragrans*, and *H. portoricensis* which were below unity, and only in one species (*Pachycereus pecten-aboriginum*) was it slightly larger than unity. The inclusion of a seed bank (WSB) increased *λ* across eleven species (*C. macrocephalus, C. mezcalaensis, C. polylophus, C. tetetzo, E. chiotilla, H. fragrans, H. portoricensis, P. gaumeri, P. pecten-aboriginum, P. pringlei*, and *S. eruca*), where values (and confidence intervals) were larger than unity (>2.0 % increase of *λ*) and *C. gigantea* that was originally below unity (Table 2). Including a hypothetical seed bank (WSB) yielded systematic significant increases in *λ* (*t*-test = 4.4784, *df* = 11, *P* = 0.001).

### 3.2 The importance of the seed bank

The PCA showed that four components accounted for 92.25% of the total variance. PC-1 explained 33.98% of the total variance with two vital rates: positive growth (*γ*) and individual fecundity (*ϕ*) with high loadings. PC-2 explained 27.10% of the residual variance with positively correlated retrogression (*ρ*) and negatively correlated with species. PC-3 explained 18.86% of the residual variance due to the presence/absence of a seed bank. PC-4 explained 12.21% of the residual variance and positively correlated with survival (*σ*) (Supplementary Appendix B). In all but PC3 matrix dimensions had no significant effects (*F*_5,66_= 0.84; P=0.52, *F*_5,66_= 0.76; P=0.58, *F*_5,66_= 1.49; P=0.20). For PC3, there was a significant difference given by the presence of a seed bank (Figure 2) regardless of matrix dimension (*F*_5,66_= 41.44; P=0.0001; Figure 3).

**FIGURE 2.**
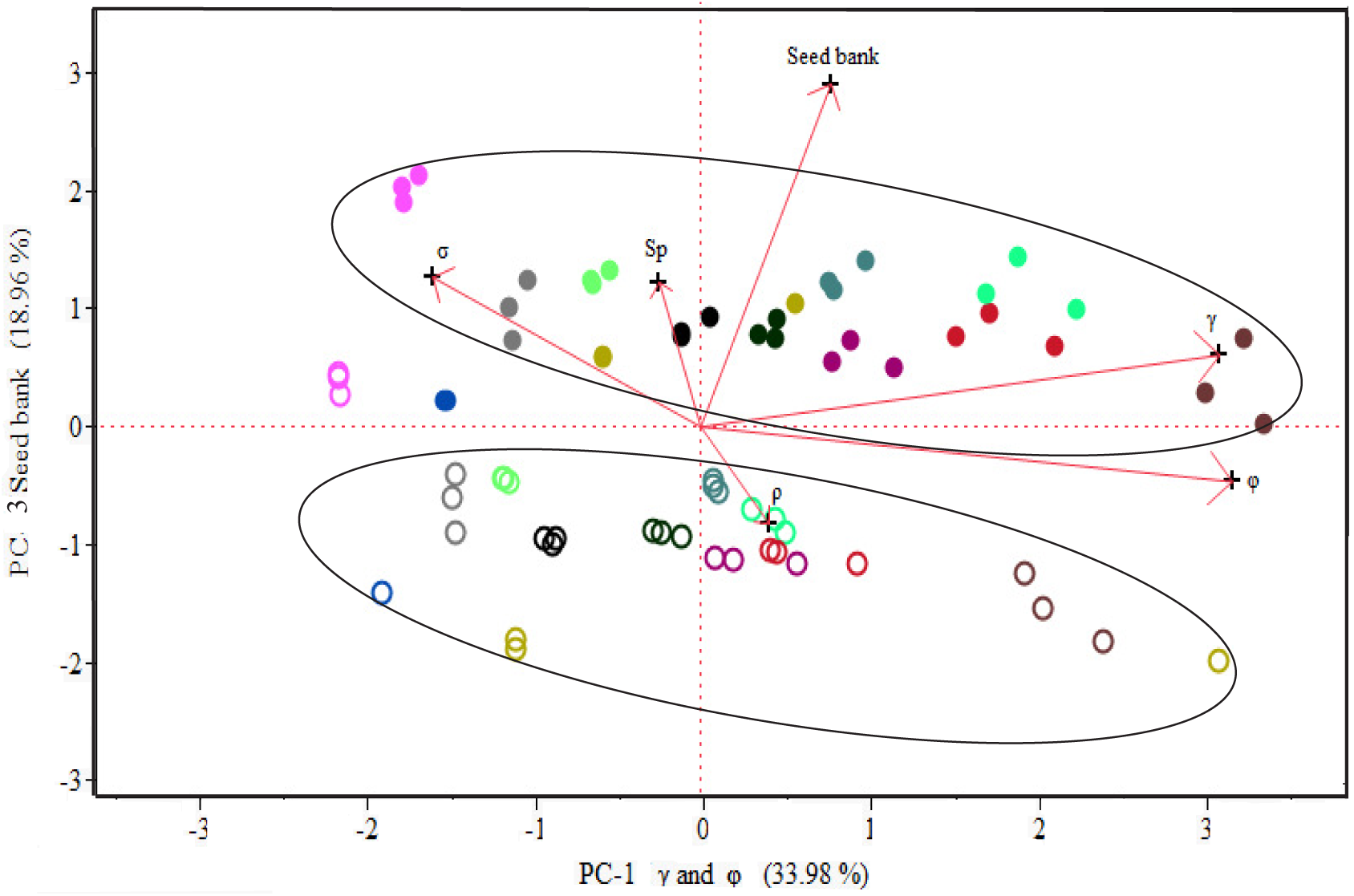
Biplot resulting from principal components analysis (PC 1-3) was used to evaluate elasticities of vital rates: survival in a given stage class (*σ*), negative growth (*ρ*), recruitment or positive growth (*γ*), individual fecundity (*ϕ*), seed bank models, and species effect (Sp). Different colors showed each species of 12 columnar cacti and each ellipse clustered two groups: WOSB (open circles) and WSB models (solid circle).

**FIGURE 3.**
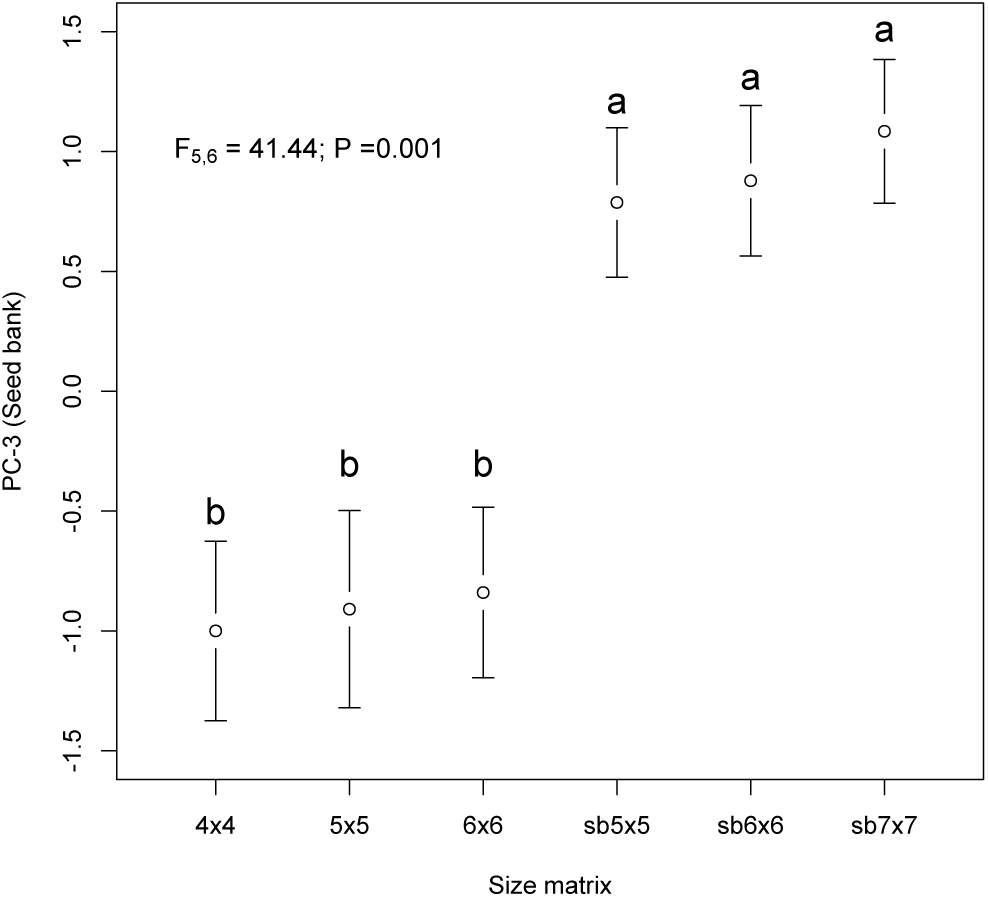
Average (± 95 CI) of PC-3 (seed bank model) against matrix dimension (WOSB = 6×6, 5×5 and 4×4; WSB = sb7×7, sb6×6, and sb5×5). Different letters indicate differences between matrix dimensions with and without seed bank (P<0.05).

### 3.3 Numerical simulations

The numerical simulations of the vital rates showed a significant effect on *λ* by seedling survival (*σ*_*se*_) (Figure 4 a and f) in both SWB and WOSB models, as well as in the seed bank (*σ*_*sb*_) (Figure 4 d) for the WSB model. Although survival in the seed bank is unknown under field conditions, simulations suggest *λ* changes significantly, even with a relatively small shift in the survival probability. Small changes in the seedling to juvenile (*λ*_*se*−*ju*_) and juvenile survival (*σ* _*ju*_) transitions for the WOSB (Figure 4 b and c) and the seed to seedling (*γ*_*sb*−*se*_) in WSB model (Figure 4 e) were particularly important. The recruitment of seedlings and their survival of the seed bank seems to be crucial processes for the population dynamics of columnar cacti except in (*C. gigantea*) and (*H. fragrans*), where changes transition from seed to seedling had a negligible impact on *λ*.

**FIGURE 4.**
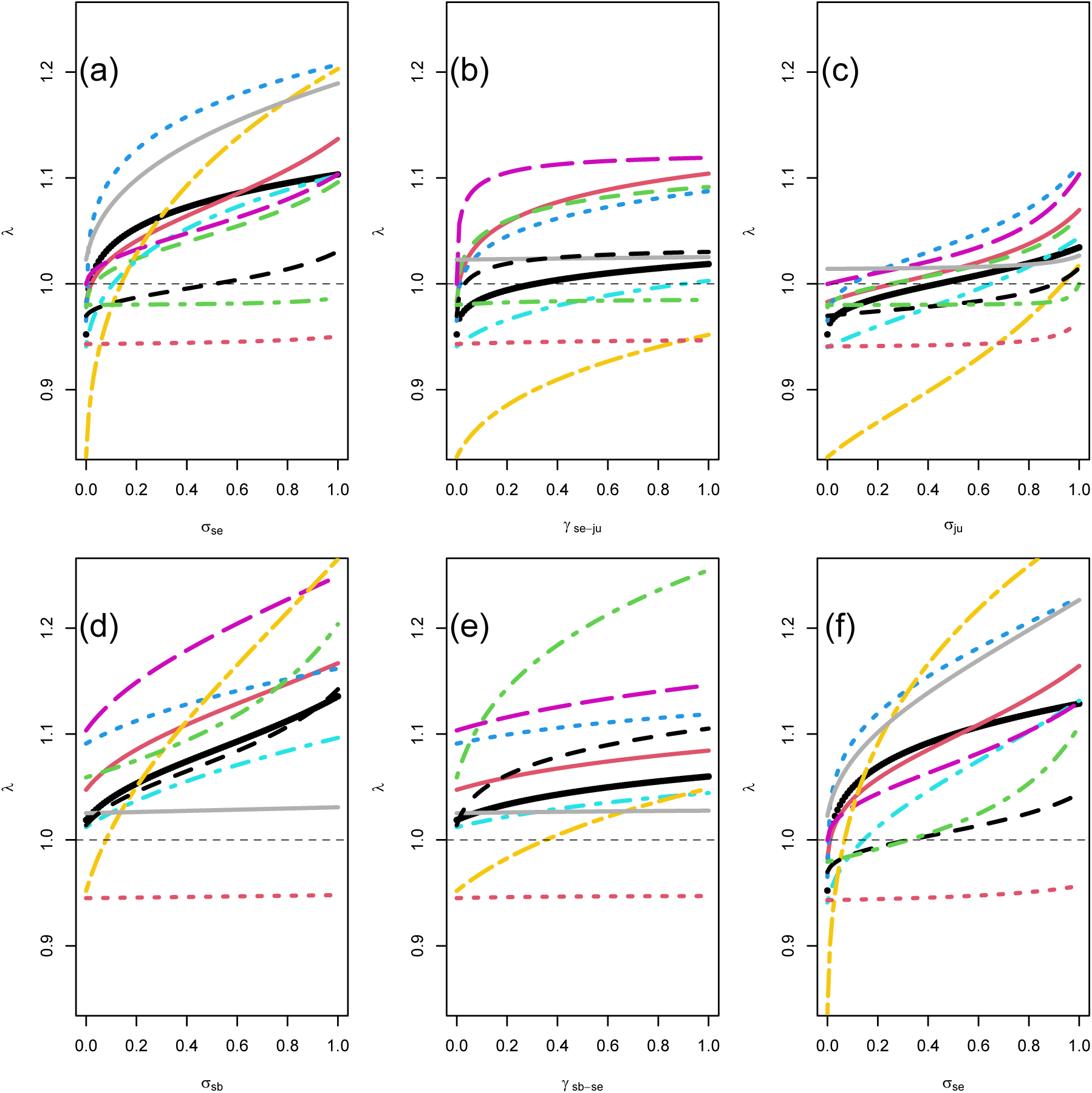
Population growth rates (*λ*) as a function of seedling survival (*σ*_*se*_) (a), seedling to juvenile (*γ*_*se*−*ju*_) (b), juvenile survival (*σ*_*ju*_) (c), seed bank (*σ* _*sb*_) (d), germination (*γ* _*sb*−*se*_) (e) and seedling survival ((*σ*_*se*_) (f). Eleven columnar cacti were using the WOSB model 6×6 (a-c) and WSB model 7×7 (d-f) and simulated by changing the values, between 0 and 1 at intervals of 0.01, from each vital rates. The grey dotted line is equilibrium, *λ* = 1. The black lines correspond to columnar cacti. It was not included *Carnegiea gigantea* to present small values of *λ* ≈ 0.50. Different line colors showed of each species of 11 columnar cacti.

## 4 DISCUSSION

Not explicitly including a seed bank in demographic models continues to be a confounding factor in understanding and modeling population dynamics in columnar cacti. There is very little information in arid environments about what is believed to be the most limiting factor for population dynamics: seed banks and seedling dynamics. This is surprising as several studies have highlighted the importance of seed banks for the persistence of populations over time in unpredictable environments (Gutterman and Venable, 2014) and the extremely limiting conditions for seedlings imposed by abiotic and biotic conditions. Even though there is growing evidence that seed banks can be found in cacti, they have largely been overlooked despite the importance towards population dynamics, especially so in the early life stages. No explicit consideration of the seed bank in a population can generate fluctuating degrees of uncertainty in the estimation of growth rates and the accuracy of the estimated vital rates (Nguyen et al., 2019).

Most studies of columnar cacti have values of *λ* that are not significantly different from equilibrium (Godínez-Álvarez et al., 2003), with relatively large confidence intervals, suggesting that populations of these species are either stable or close to equilibrium (Rae and Ebert, 2002; Méndez et al., 2004; Morales-Romero et al., 2012). Unfortunately, confidence intervals are sufficiently large that any management decision should be taken with caution if at all. The decrease of *λ* in two species (*C. gigantea*, and *H. fragrans*) may be caused by species or even population-specific factors and inter-annual variations in climatic factors. External factors are commonly determinants of the endangered status for cacti species (Goettsch et al., 2015) including some columnar species (*Carnegiea gigantea, P. gaumeri, S. eruca, H. fragrans* and *H. potoricensis*). The drivers of declining populations are usually associated to fragmentation and habitat loss (urbanization, road construction, cattle ranch management and agriculture, Esparza-Olguín et al., 2002; Méndez et al., 2004; Rojas-Sandoval and Meléndez-Ackerman, 2013) as well as interannual variation in climatic factors (Esparza-Olguín et al., 2002, 2005; Arroyo-Cosultchi et al., 2016).

Overall, the phenomenon of *λ* close to unity is not surprising and is actually expected for long-lived species such as cacti, in which relevant population processes may occur at the scale of decades (Pierson et al., 2013), slow growth, late maturity, low fecundity, and high survival probabilities are common life-history traits (Esparza-Olguín et al., 2002; Godínez-Álvarez et al., 2003). Results in this study indicate that columnar species of cacti are at equilibrium with structures mainly composed of juvenile and young adults and consistent low numbers of seedling numbers (except for *Cephalocereus polylophus* (Arroyo-Cosultchi et al., 2016) *and C. tetetzo* (Godínez-Alvarez and Valiente-Banuet, 2004)). A limitation of this study was the minimization of interannual and interpopulation variability by averaging matrices as well as ignoring episodic interannual recruitment, although these were out of the scope of our research.

An increase in *λ* followed the addition of a hypothetical transient seed bank. Seed banks seem to be more widespread than previously thought as evidence suggests short term seed banks in the subtribe Stenocerinae (*Myrtillocactus geometrizans, Polaskia chende, Stenocereus* sp. and *Stenocereus stelatus*) (Ordoñez, 2008; Cano-Salgado et al., 2012; Álvarez-Espino et al., 2014; Ordoñez Salanueva et al., 2017) and the tribe Trichocereeae (*Harrisia fragrans*) (Goodman et al., 2012). Even though adult longevity in the Cactaceae is high and seed banks would not be theoretically expected, seed banks decouple reproduction from other life stages which buffer against environmental variation. A clearer understanding of age-dependent germination rates of seeds, age-dependent survival of non-germinated seeds, and the production of new seeds by reproductive plants (Doak et al., 2002) is needed to determine the specific factors (e.g., environmental, physiological) that contribute to their formation. The presence of seed banks changes the life history of species and has a small but positive consequence of population growth rates that may compound population dynamics in variable environments.

The population dynamics of the majority of the studied species strongly depends on the survival of adult individuals and the growth of intermediate stages in the life cycle, a pattern similar to that reported for many long-lived plants including succulents, shrubs, and trees (Silvertown et al., 1993; Enright et al., 1995; Franco and Silvertown, 2004). Arid and semi-arid environments pose important challenges for plant persistence, and species rely on recruitment whereby the lack of recruitment at any given time gives the impression of a slowly decreasing population that depends on survival (Holland and Molina-Freaner, 2013). In the case of *C. gigantea, H. portoricensis* and *C. polylophus*, the recurring presence of freezing and ENSO have been shown to be phenomena that strongly impacted populations by either high mortality or recruitment (Pierson et al., 2013; Rojas-Sandoval and Meléndez-Ackerman, 2013; Arroyo-Cosultchi et al., 2016). The lack of recruitment in the studies on columnar and other cacti species points towards a limiting demographic stage, and has often been associated to seed predation (seed limitation) and/or high seedling mortality (seedling limitation) (Mandujano et al., 2001; Esparza-Olguín et al., 2002, 2005; Ferrer-Cervantes et al., 2012; Rojas-Sandoval and Meléndez-Ackerman, 2013; Zepeda-Martínez et al., 2013).

Low water availability and the quantity of solar radiation that characterize arid and semi-arid environments impose serious limitations on population growth, mainly because they induce high seedling mortality and limit the establishment of new individuals (Steenbergh and Lowe, 1977). The PCA allowed us to identify that the vital rates corresponding to positive growth and fecundity were higher, so these vital rates had significant effects on population dynamics. The results of the analysis variance of the seed bank inclusion were shown important and the effect of the matrix size was negligible. Adding a seed bank increased the importance of the vital rates (positive growth and fecundity) for the early life stages and our results from numerical simulations showed that changes in seedling survival and seed bank could have significant effects on the population dynamics of columnar cacti and therefore protecting the seed bank is essential to the persistence of theses species.

The simulations suggest that in most cases, seedling limitation has a larger effect than seed limitation in the population dynamics of cacti. Columnar cacti are strongly seed-limited by the variable reproduction of adults, and the high predation of seed but are also seedling-limited as even when enough seeds are produced, seedlings do not survive. The early stages are a possible option for the management of cacti species which should consider manipulations to enhance/reduce recruitment by either active introduction/elimination of juveniles or by increasing/decreasing the survival probabilities of naturally established plants. For example, several species of *Harrisia* (*H. balansae, H. martinii, H. pomanensis* and *H. tortuosa*) are considered highly invasive (Novoa et al., 2015) and control of early stages could help manage these populations. On the other hand, four species (*H. fragrans, H. portoricensis, P gaumeri* and *C. polylophus*) have some degree of endangered status so increasing the seed-seedling transitions can provide solutions for conservation strategies. The cost-efficient management of cacti would indicate that for conservation purposes, the reintroduction by transplanting nursery reared seedlings or juveniles (reducing seedling limitation) to be a better strategy than sowing seeds (seed augmentation) directly into the wild (Birnbaum et al., 2011; Reemts et al., 2014).

Seed banking may increase seedling recruitment of columnar cacti by increasing seedling opportunities when conditions are favorable for survival. Their effect would also suggest that several columnar cacti populations are not threatened in demographic conditions under the assumptions used in this study. This by no means suggests that other contributing factors to their decline should not be considered for conservation (see Goettsch et al., 2015). If we consider that Rojas-Aréchiga (2014) found positive photoblastism and seed size as phylogenetically associated to the subtribe Stenocerinae, at least the physiological component of seed bank formation is favored in this group (Rojas-Aréchiga and Batis, 2001). We are in need to clearly determine the presence and longevity of seeds for many cacti species and untangle the factors behind seed and seedling limitation to adequately portray the life cycle of this taxonomic group.

## Acknowledgements

This research is part of the doctoral studies of Gabriel Arroyo-Cosultchi (UAM-X). Financial support was provided by CONACyT (165908) to MCM, CONACyT sabbatical leave scholarship to JG and PASPA-DGAPA sabbatical scholarship to MCM. Comments by M. Rojas Aréchiga and M. Franco significantly improved this manuscript. Thanks are also due to M. Franco and R. Clark-Tapia, who made available the demographic data for *Carnegiea giganteae* and *Stenocereus eruca*.

## Appendix Supplementary data

Supplementary material related to this article can be found, in the online version, at doi:.

